# Steric accessibility of the *N*-terminus improves the titer and quality of recombinant proteins secreted from *Komagataella phaffii*

**DOI:** 10.1101/2022.06.17.496632

**Authors:** Neil C. Dalvie, Christopher A. Naranjo, Sergio A. Rodriguez-Aponte, Ryan S. Johnston, J. Christopher Love

## Abstract

**Background:** *Komagataella phaffii* is a commonly used alternative host for manufacturing therapeutic proteins, in part because of its ability to secrete recombinant proteins into the extracellular space. Incorrect processing of secreted proteins by cells can, however, cause non-functional product-related variants, which are expensive to remove in purification and lower overall process yields. The secretion signal peptide, attached to the *N*-terminus of the recombinant protein, is a major determinant of the quality of the protein sequence and yield. In *K. phaffii*, the signal peptide from the *Saccharomyces cerevisiae* alpha mating factor often yields the highest secreted titer of recombinant proteins, but the quality of secreted protein can vary highly.

**Results:** We determined that an aggregated product-related variant of the SARS-CoV-2 receptor binding domain is caused by *N*-terminal extension from incomplete cleavage of the signal peptide. We eliminated this variant and improved secreted protein titer up to 76% by extension of the *N*-terminus with a short, functional peptide moiety or with the EAEA residues from the native signal peptide. We then applied this strategy to three other recombinant subunit vaccine antigens and observed consistent elimination of the same aggregated product-related variant. Finally, we demonstrated that this benefit in quality and secreted titer can be achieved with addition of a single amino acid to the *N*-terminus of the recombinant protein.

**Conclusions:** Our observations suggest that steric hindrance of proteases in the Golgi that cleave the signal peptide can cause unwanted *N*-terminal extension and related product variants. We demonstrated that this phenomenon occurs for multiple recombinant proteins, and can be addressed by minimal modification of the *N*-terminus to improve steric accessibility. This strategy will enable consistent secretion of a broad range of recombinant proteins with the highly productive alpha mating factor secretion signal peptide.

## Background

Secreting heterologous recombinant proteins into the extracellular medium during fermentation of microbial organisms can intensify processes for production, and in turn reduce operational costs and complexities [1]. Secretion of proteins by cells also enables alternative operating modes like perfusion for continuous processing, and simplifies subsequent recovery of the protein from culture supernatant instead of cellular lysates [2–4].

Canonical cellular processes for secreting proteins in eukaryotic microorganisms requires translocation of the translated protein. An *N*-terminal polypeptide sequence directs translocation of the protein to the endoplasmic reticulum (ER) from which the protein is ultimately transported via vesicles through the Golgi and to the cell’s surface [5]. The nature of these ‘signal’ peptides, along with the features of the heterologous protein itself, can vary the production yields, primary, secondary, or tertiary structures, and post-translational modifications. Proteins with *N*-terminal truncations or extensions, proteolytic cleavage, or aberrant N-or O-linked glycosylation may be non-functional, immunogenic, prone to aggregation, or unstable in formulation [6]. These product-related variants are often challenging to remove by affinity chromatography and typically require additional process operations like ion exchange or hydrophobic interaction chromatography [7–9]. To maximize the benefits of continuous production of proteins by secretion and subsequent intensification in recovery, it is essential to minimize heterogeneity in the proteins produced by the host cells.

The yeast *Komagataella phaffii* (*Pichia pastoris*) is a common alternative host with high potential for low-cost manufacturing of therapeutic proteins like vaccine antigens and monoclonal antibodies [10,11]. It has a highly developed secretory pathway, grows quickly to high cell densities, and can be easily genetically modified [12–14]. The most commonly used signal peptide to direct secretion of recombinant proteins in *K. phaffii* is the α-mating factor signal peptide (αSP) from *Saccharomyces cerevisiae*, which comprises a 19 amino acid [pre] sequence and a 67 amino acid [pro] sequence, which terminates in a Glu-Ala-Glu-Ala (EAEA) motif [15,16]. In *S. cerevisiae*, the [pre] sequence is removed in the endoplasmic reticulum, the [pro] sequence, which includes multiple sites for *N*-glycosylation, is removed by the KEX2 protease in the Golgi; the STE13 protease removes the residual EAEA amino acids at the N-terminus of the secreted α-mating factor [17].

In *K. phaffii*, however, efficient processing of the αSP can depend on the recombinant heterologous protein used. Improper or inefficient cleavage of the signal peptide can impact the secreted titer of the correct recombinant protein or create product variants like *N*-terminal extension or truncation [18–21]. Secretion of recombinant human interferon-α2b, for example, resulted in incomplete removal of the *N*-terminal EAEA residues [22]. In this case, the KEX2 cleavage site was sufficient to cleave the [pro] signal peptide when EAEA was removed from the coding sequence [3]. Indeed, the EAEA residues have often been omitted from the αSP when used for secretion of recombinant proteins from *K. phaffii* [19,23,24]. Despite inconsistent *N*-terminal processing and design practices, the *S. cerevisiae* αSP remains the signal peptide most commonly used in *K. phaffii* because very few alternative signal peptides have resulted in consistently higher product quality or secreted titers of recombinant protein [24–26]. Refined understanding of the requirements for efficient processing of the aSP in *K. phaffii* would, therefore, enhance designs to improve protein secretion in this host.

Here, we evaluated the impact of signal peptide processing on the quality and secreted titer of several recombinant therapeutic proteins, and tested approaches to minimize variations by altering the EAEA motif. We observed that an aggregated product-related variant of the SARS-CoV-2 receptor binding domain (RBD) can result from *N*-terminal extension from incomplete processing of the signal peptide. We eliminated this product variant and increased secreted titers by steric extension of the *N*-terminus of the RBD first by a functional peptide and the αSP-EAEA residues, and then tested whether or not similar benefits could be obtained using only single amino acids. Finally, we demonstrated that extension of the *N*-terminus of three other subunit vaccine antigens by αSP-EAEA residues or a peptide epitope also eliminates an aggregated product-related variant. Together, these results provide evidence from multiple examples of recombinant proteins for how inefficient cleavage of the signal peptide can lead to variations of different quality attributes of the proteins, and suggest some strategies for improving protein quality with minimal sequence modifications.

## Results

### Incomplete cleavage by KEX2 leads to protein aggregation

High-molecular weight species or aggregated recombinant proteins can impact final yields and may be hard to remove since they may have similar biophysical features as properly folded protein [27]. We previously reported manufacturing of the SARS-CoV-2 receptor binding domain (RBD), a promising candidate subunit vaccine antigen for COVID-19, in *K. phaffii* [28,29]. We observed a high-molecular weight species (RBD-HMW) in purified RBD samples purified by ion exchange chromatography. We developed one strategy to reduce these species by engineering the RBD sequence to reduce the intrinsic aggregation of the molecule [30]. Nonetheless, we still observed RBD-HMW after purification with this engineered version.

We therefore sought to further investigate features of the RBD-HMW. We separated the RBD-HMW from monomeric RBD by SEC (Fig. 1A), and treated the RBD-HMW with PNGase. This treatment showed the RBD-HMW (∼70-100 apparent kDa) comprised several distinct species of approximately ∼30 kD (Fig. 1B). These data showed the formation of the RBD-HMW species depends on N-glycosylation. We next analyzed the intact mass of these species by liquid chromatography mass spectrometry (LCMS) and observed that these polypeptides retained portions of the αSP (Fig. 1C). These included residual deglycosylated fragments of the [pro] region of the αSP, ranging from 9 to 66 amino acids (Table 1). While the recombinant RBD molecule contains only one N-linked glycosylation site (N12), the fragment of the [pro] peptide appended to the *N*-terminus contains three additional canonical sites for N-linked glycosylation. The αSP also contains a predicted sites for O-linked glycosylation (T25, NetOGlyc 4.0), and we observed O-glycosylation of the *N*-terminal extensions (Fig. 1C).

**Table 1.**
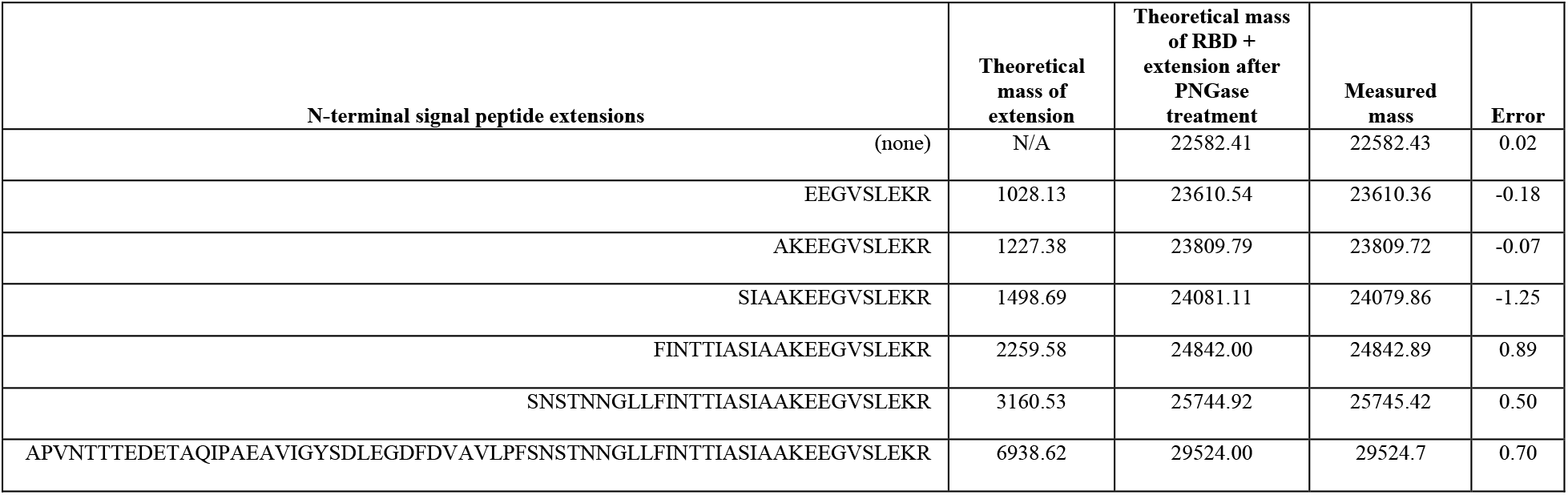
*N*-terminal extensions of the RBD (amu).

**Fig. 1.**
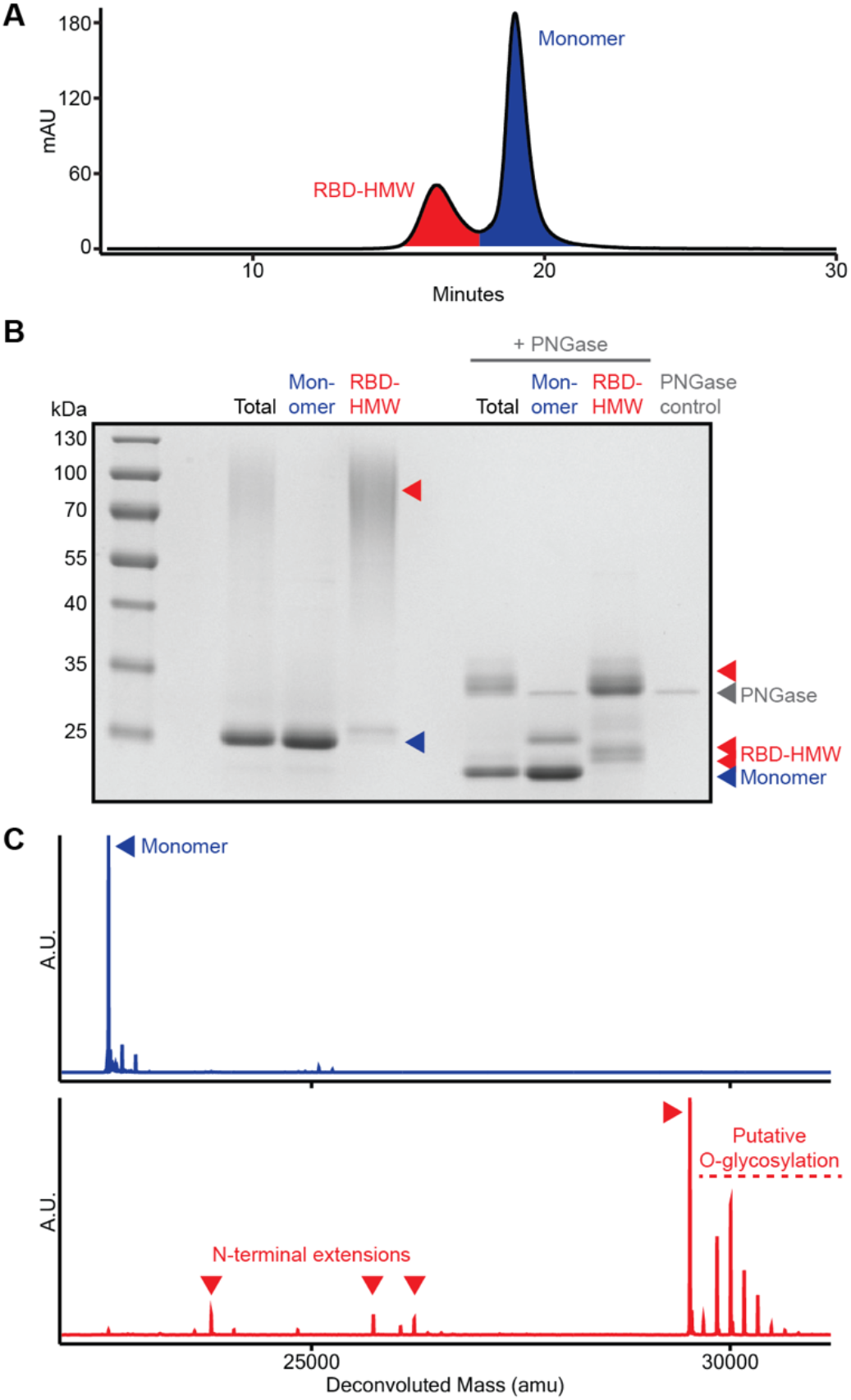
Aggregated product-related variant is caused by *N*-terminal extension. A) Separation of the aggregated RBD variant from monomeric RBD by size exclusion chromatography. B) Reduced SDS-PAGE of purified RBD and each fraction after separation by size exclusion chromatography. C) Intact LCMS of each RBD fraction.

### N-terminal modification eliminates N-terminal extension

Next, we investigated if *N*-terminal extensions were evident in other engineered variations of the RBD. Specifically, we assessed the quality of RBD antigens expressed in *K. phaffii* that were genetically modified to include a 13 amino acid SpyTag peptide, a motif for transpeptidation useful to link proteins onto other proteins modified with SpyCatcher, such as protein nanoparticles [31,32]. We appended the SpyTag peptide to either the *C*-terminus (RBD-SpyTag) or the *N*-terminus (SpyTag-RBD) of the RBD antigen with a flexible linker sequence GGDGGDGGDGG. Interestingly, we observed the RBD-HMW only in purified RBD-SpyTag (Fig. 2A). We confirmed this observation by LCMS, and the observed *N*-terminal extensions similar to the unmodified RBD (Fig. 2B). SpyTag-RBD, on the other hand, exhibited no detectable *N*-terminal extensions, suggesting that the endogenous KEX2 protease fully processes the [pro] peptide when fused to SpyTag-RBD. We reasoned that protrusion of the SpyTag peptide from the *N*-terminus of the folded RBD molecule may allow KEX2 protease improved steric access to the dibasic cleavage motif (KR). Indeed, inspection of the predicted structure of the SpyTag-RBD suggested that the *N*-terminus of SpyTag-RBD may be more exposed than the *N*-terminus of unmodified RBD (Fig. S1).

**Fig. 2.**
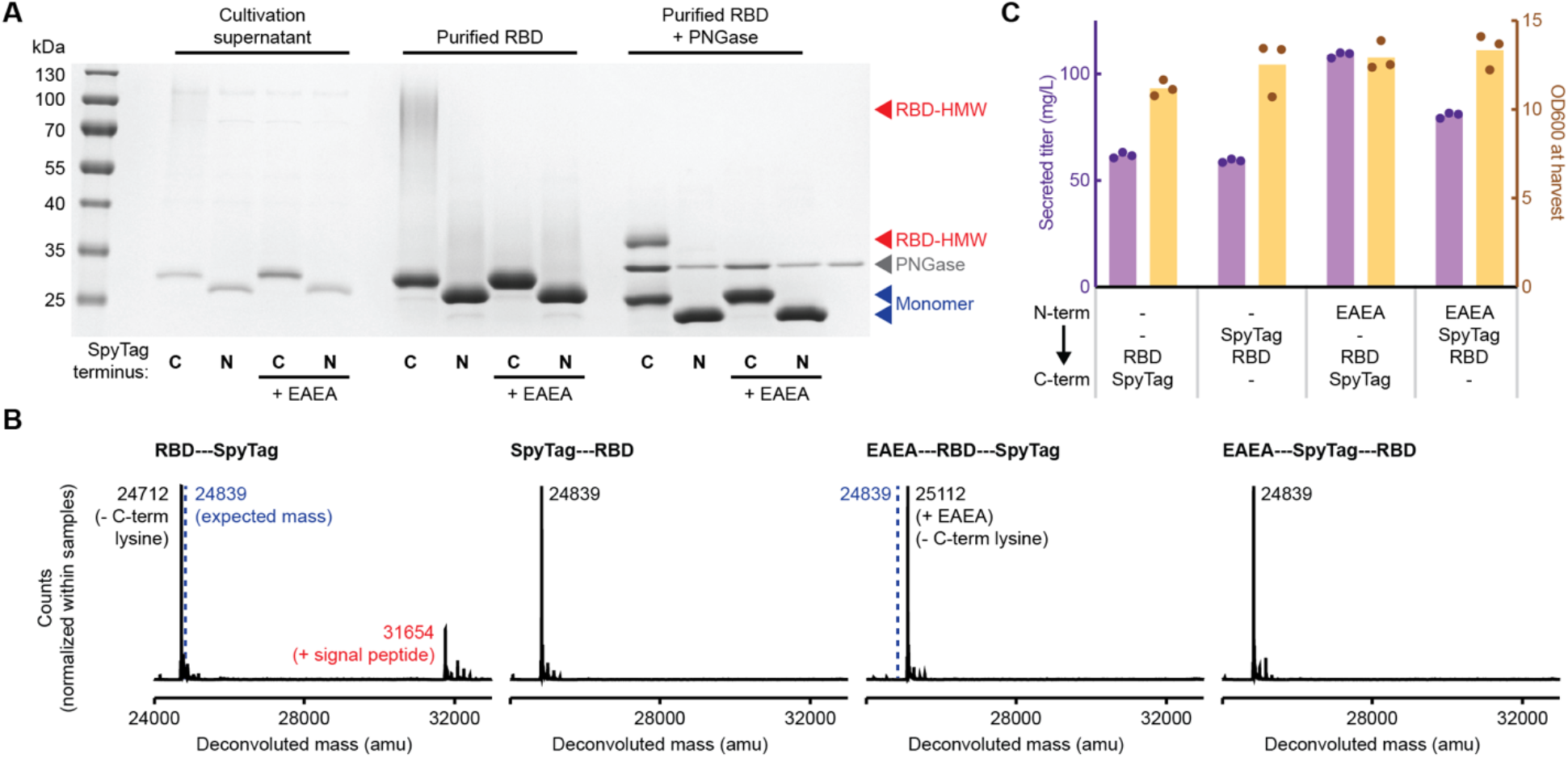
Addition of residues to the *N*-terminus eliminates *N*-terminal extension. A) Reduced SDS-PAGE of unpurified supernatant and purified variants of the RBD. B) Intact LCMS of purified RBD variants. C) Secreted titer and cell density of strains producing RBD variants with different *N*-terminal sequences. Cells were cultivated at 3 mL scale for one day of outgrowth and one day of production. Bars represent the average of three independent cultures.

Based on our observations with SpyTag-RBD, we hypothesized that complete cleavage of the signal sequence by KEX2 could eliminate *N*-terminal extension and the RBD-HMW. To facilitate cleavage by KEX2, we re-inserted the EAEA motif natively found in the *S. cerevisiae* αSP cleavage site (KR-EAEA) to the *N*-terminus of both RBD-SpyTag and SpyTag-RBD. As expected, addition of EAEA eliminated the RBD-HMW that was previously observed in purified RBD-SpyTag (Fig. 2A). We also performed LCMS to confirm the absence of *N*-terminal extensions for both antigens expressed with EAEA (Fig. 2B). Interestingly, the EAEA residues remained at the N-terminus of the RBD-SpyTag antigen, but were removed from the SpyTag-RBD antigen. During processing of the native αSP in *S. cerevisiae*, these residues are removed by the protease STE13 [15]. We hypothesized that the protrusion of the *N*-terminus of the protein afforded by the SpyTag peptide reduced steric interference from the folded RBD itself, allowing efficient access and cleavage by the STE13 protease in *K. phaffii*. Of note, we also consistently observed removal of the *C*-terminal lysine residue for antigens with SpyTag appended at the *C*-terminus (Fig. 2B). In our experience, however, this variant does not impact the efficiency of conjugation to SpyCatcher or the performance of RBD-SpyTag as a vaccine antigen [30].

In addition to assessing the impact of SpyTag and EAEA residues on product quality, we also assessed the yield of the antigens produced by *K. phaffii*. We observed an increase in secreted titer of the RBD antigens modified with EAEA residues (Fig. 2C). We observed the largest titer improvement from 62 mg/L to 109 mg/L (∼76% increase) upon adding EAEA residues to the RBD-SpyTag, which also eliminated the RBD-HMW. We previously hypothesized that RBD antigens that tend to aggregate after purification may also aggregate inside the *K. phaffii* secretory pathway (and in turn trigger a secretory stress response and lower the secreted titer) [30]. Interestingly, we also observed an improvement in secreted titer from 59 mg/L to 81 mg/L (∼36%)when we added EAEA residues to the SpyTag-RBD, despite observing no *N*-terminal extension and no aggregation of the original SpyTag-RBD. This observation suggests that, in addition to reducing product-related variants, efficient cleavage of the αSP by KEX2 may generally enhance the secretion of recombinant proteins from *K. phaffii*.

### N-terminal engineering of rotavirus subunit antigens

Next, we sought to determine whether or not these findings were specific only to the RBD. We previously reported the manufacturing of an engineered trivalent subunit vaccine for rotavirus in *K. phaffii* [18]. In this vaccine design, each of the three truncated VP8 antigens (P[4], P[6], and P[8]) was genetically linked to a tetanus toxoid epitope (P2) to enhance the immunogenicity of the vaccine [33]. We hypothesized that P2, which is 15 amino acids and attached by a small flexible linker, would enable efficient cleavage of the αSP by KEX2 protease from P[4], P[6], and P[8],similar to the effect observed with the SpyTag peptide for the SARS-CoV-2 RBD. To test this hypothesis, we expressed each engineered VP8 antigen with and without the P2 epitope, and with and without the αSP EAEA residues (Fig. 3A). For the P[4] and P[8] antigens, we observed that the addition of the P2 epitope, the αSP EAEA residues, or both, to the *N*-terminus of the proteins eliminated HMW species evident in culture supernatants; in contrast, unmodified antigens tended to form HMW species. In our previous work, we found P[6] antigen secreted at 4-10x lower titers than P[4] and P[8] [18]. In this study, appending the P2 epitope, the αSP EAEA residues, or both to the *N*-terminus did not substantially improve the titers of the P[6] antigen, but there was no detectable P[6] antigen when there was no *N*-terminal modification. We hypothesize that the unmodified P[6] may also form HMW species, but that the HMW species was not present at a high enough concentration for detection by SDS-PAGE. These data also further corroborate that intrinsic differences in the sequence of the P[6] antigen itself contribute to the poor titers observed relative to the other two serotypes tested here [18].

**Fig. 3.**
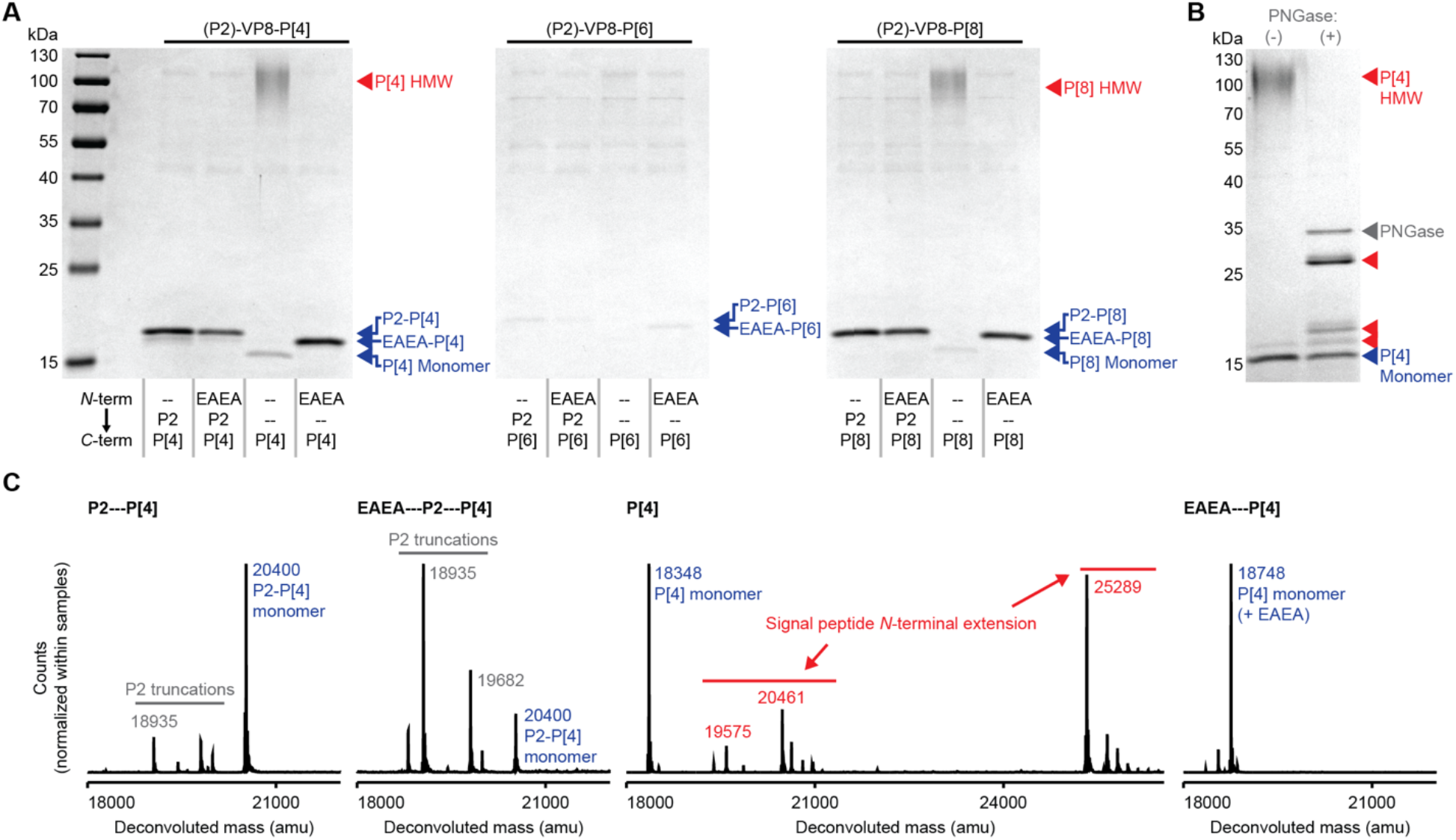
*N*-terminal modification of rotavirus VP8 antigens. A) Reduced SDS-PAGE of unpurified supernatant of variants of P[4], P[6], and P[8] antigens. B) Reduced SDS-PAGE of purified P[4] with no *N*-terminal modification. C) Intact LCMS of purified P[4] variants.

To determine if the HMW variant of the rotavirus VP8 subunits was similar to the HMW variant of the SARS-CoV-2 RBD, we evaluated the primary structure of the four versions of the P[4] antigen (P2-P[4], EAEA-P2-P[4], P[4], and EAEA-P[4]). We cultivated each strain in shake flasks and purified the secreted antigens. First, we treated the P[4]-HMW protein with PNGase, and observed several distinct HMW species, similar to the RBD-HMW (Fig. 3B). Next, we performed intact LCMS on each P[4] variant (Fig. 3C). We observed that the P[4]-HMW species from the unmodified P[4] antigen comprised *N*-terminal extensions, which is consistent with our previous observations of the RBD-HMW. Over 70% of the purified P[4] comprised P[4]-HMW species by LCMS. We noticed, additionally, that EAEA residues remained when appended to the *N*-terminus of the P[4] antigen, but were removed by STE13 protease when appended to the *N*-terminus of the P2 epitope. Indeed, the apparent molecular weights of P2-P[4/6/8] and EAEA-P2-P[4/6/8] observed by SDS-PAGE were the same for all three antigens (Fig. 3A). Finally, we observed *N*-terminal truncation of the P2 epitope itself (Fig. 3C). We previously have observed these truncated variants of P2-P[4], and we hypothesized that this truncation is mediated by a native serine protease from *K. phaffii*. Interestingly, this observation is also consistent with our hypothesis here: that reducing steric hinderance at the *N*-terminus of the P2 epitope improves its accessibility and consequently renders higher protease activity. Indeed, we did not observe any *N*-terminal truncation of the EAEA-P[4] antigen, which lacks the P2 epitope. Together, these results suggest that protrusion of the *N*-terminus away from the folded structure of the protein facilitates efficient protease activity, reducing aberrant *N*-terminal extension and aggregation of multiple recombinant secreted proteins.

### Elimination of aberrant N-terminal extension with minimal N-terminal engineering

Throughout this study, we observed that the EAEA residues often remained on the secreted protein when placed directly adjacent to the globular RBD or VP8 proteins (Fig. 2B, 3B). For some vaccine antigens, EAEA residues may be an acceptable modification to the sequence of a candidate drug substance, given the large improvements in both the quality and titer of the secreted product. EAEA could, however, present potential risks or perceived concerns of unwanted immunogenicity for other parenteral therapeutic proteins. We sought, therefore, to determine the minimal number of residues necessary to eliminate *N*-terminal variants containing a full or partial signal peptide.

We expressed the unmodified RBD antigen using the αSP with different numbers of EA repeats as well as different single amino acids at the *N*-terminus. We observed that addition of a single EA repeat or a single E residue conferred a similar increase in the secreted titer of the RBD compared to the unmodified RBD antigen (Fig. 4A). Additional EA repeats did not appear to further improve the titer or quality of the RBD antigen. Interestingly, we observed that most single amino acids were sufficient to eliminate the RBD-HMW species (Fig. 4B,C). One notable exception was proline, which appears to increase the abundance of the RBD-HMW. We hypothesize that KEX2 is unable to cleave KR-P. Some amino acids like tryptophan and cysteine did reduce the secreted titer of the RBD, which we hypothesize was due to conformational changes from the large size of tryptophan or dimerization of the RBD from addition of a free cysteine residue. Based on these observations, addition of a small, charged or polar amino acid to the *N*-terminus of the RBD appears to confer the same benefit that we observed with the EAEA residues.

**Fig. 4.**
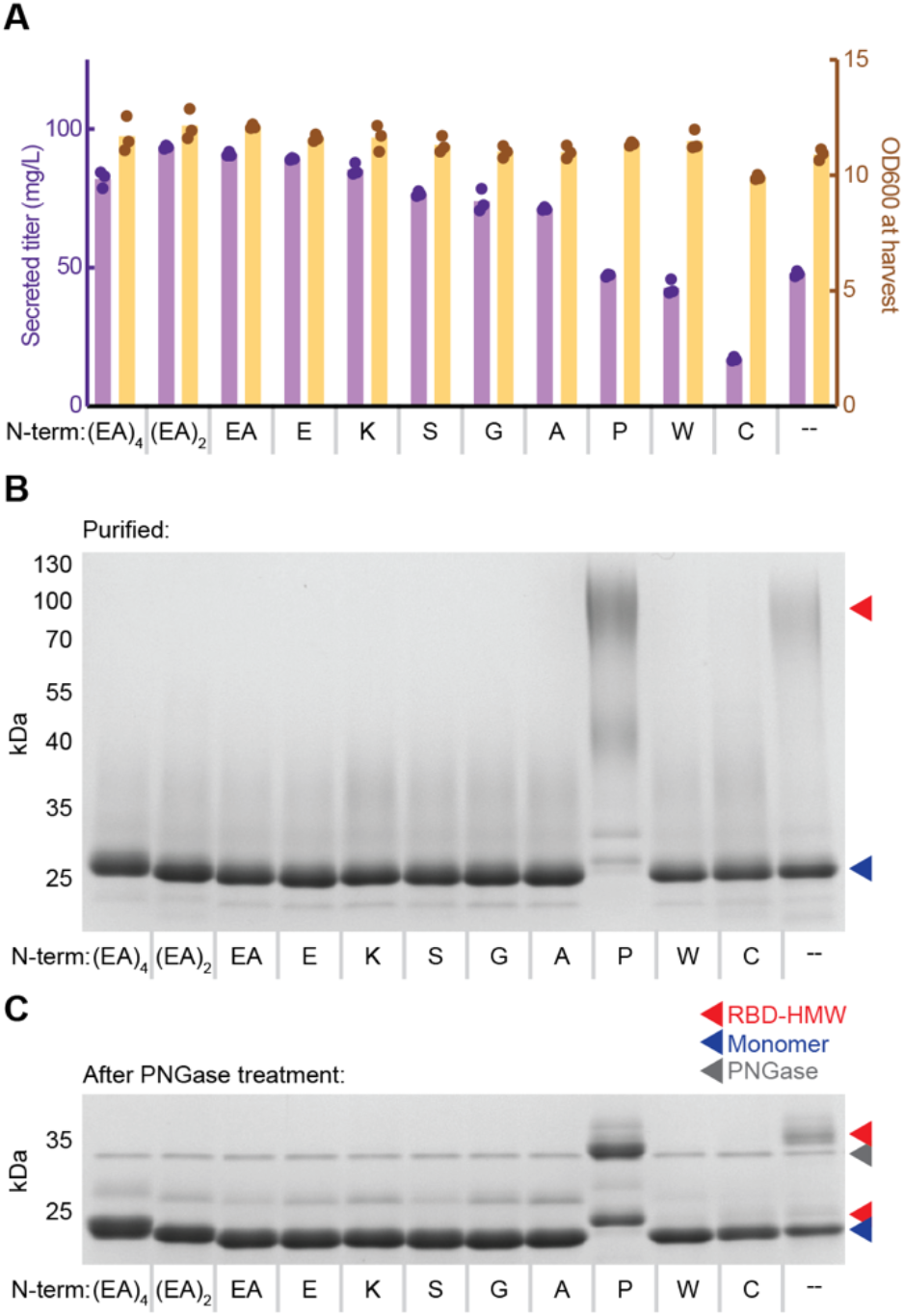
Single amino acids eliminate *N*-terminal extension. A) Secreted titer and cell density of strains producing RBD variants with different *N*-terminal sequences. Cells were cultivated at 3 mL scale for one day of outgrowth and one day of production. Bars represent the average of three independent cultures. B) Reduced SDS-PAGE of purified RBD variants. C) Reduced SDS-PAGE of purified RBD variants after treatment with PNGase.

## Discussion

We determined that a prominent product-related variant of the SARS-CoV-2 RBD secreted from yeast is caused by incomplete cleavage of the αSP. We noticed that attachment of an additional peptide sequence to the *N*-terminus of the RBD abrogated this variant, which suggested that steric accessibility of the *N*-terminus of the recombinant protein to KEX2, which cleaves the [pro] αSP in the Golgi, may determine the extent of *N*-terminal extension (Fig. 5). We also demonstrated that this phenomenon was consistent for a different set of subunit vaccine antigens, with a different peptide moiety attached to the *N*-terminus. These data together suggest that efficient cleavage of the [pro] αSP occurs when the site for cleavage in the N-terminus is not sterically occluded by globular domains.

**Fig. 5.**
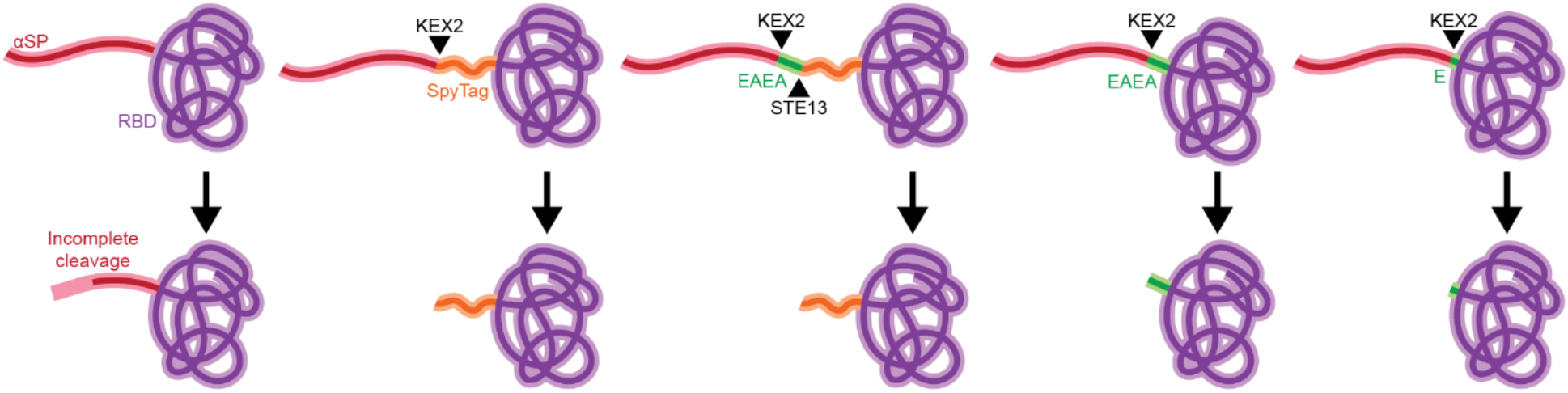
Schematic of *N*-terminal steric accessibility.

We also showed the insertion of the αSP EAEA residues can also eliminate these N-terminal variants. Others have demonstrated that mutation of the +1 residue of the KEX2 cleavage site may impact signal peptide processing and improve the secreted titer of recombinant proteins [34]. In most studies that evaluate the impact of the EAEA residues, however, the residues remain on the recombinant protein; that is, cleavage of the EAEA residues by STE13 does not occur [19,20]. These reports are consistent, therefore, with our hypothesis that protrusion of the *N*-terminus outside the folded protein domain may also facilitate cleavage by KEX2, in addition to extension of the recognized cleavage site [35]. Indeed, addition of an *N*-terminal leader sequence has been shown by others to improve secreted productivity [36].

Finally, we demonstrated that including a single amino acid to the *N*-terminus of a recombinant protein confers similar benefits in quality and secreted titer as the complete EAEA motif. Parenteral biopharmaceuticals produced in bacteria like *Escherichia coli* —including ones given for chronic indications like diabetes—include a *N*-terminal methionine that is necessary to initiate protein translation [37]. This precedent suggests that drug substances produced in *K. phaffii* could invoke a similar insertion with minimal risk or impact on clinical use. Our data here suggest further that this inclusion could provide additional benefits by minimizing N-terminal variations and enhancing titers of the heterologous protein, and that the flexibility of selecting among charged, polar, or small amino acids for the N-terminal extension could avoid other potential product-related variations like the oxidation of methionine, which is a common quality attribute measured for bacterially-expressed drug substances [38].

## Conclusions

In this study, we determined that an aggregated product-related variant of several recombinant proteins secreted from *K. phaffii* was caused by incomplete cleavage of the secretion signal peptide. We eliminated this variant by genetic addition of amino acids or short peptide moieties to the *N*-terminuses of the recombinant proteins. We hypothesize that these modifications reduce steric obstruction of the proteases KEX2 and STE13, which process the αSP. These observations suggest that steric accessibility of the *N*-terminus can determine quality and yield of a broad range of recombinant proteins produced in *K. phaffii*.

## Materials and methods

### Yeast strains

All strains were derived from wild-type *Komagataella phaffii* (NRRL Y-11430), in a modified base strain (RCR2_D196E, RVB1_K8E) described previously [39]. All RBD sequences were based on an engineered version of the RBD described previously (RBD-L452K-F490W), and were transformed as described previously [14,30]. Modifications of the RBD including EAEA, peptide moieties, and single amino acids were made by KLD site directed mutagenesis (New England Biolabs). Rotavirus antigens were designed based on optimized sequences described previously, ordered from Codex DNA, and assembled on a BioXP into the same custom vector as the RBD [18]. All plasmid sequences are found in the Supplemental Information.

### Cultivations

Strains for initial characterization and titer measurement were grown in 3 mL culture in 24-well deep well plates (25°C, 600 rpm), and strains for protein purification were grown in 100 mL culture in 500 mL shake flasks (25°C, 300 rpm). Cells were cultivated in complex media (potassium phosphate buffer pH 6.5, 1.34% nitrogen base without amino acids, 1% yeast extract, 2% peptone). Cells were inoculated at 0.1 OD600, outgrown for 24 h with 4% glycerol feed, pelleted, and resuspended in fresh media with 1% methanol and 40 g/L sorbitol to induce recombinant gene expression. Supernatant samples were collected and filtered after 24 h of production. Supernatant titers were measured by reverse phase liquid chromatography as described previously [30]. Purification of the RBD and rotavirus antigens was performed as described previously [18,30].

### Analytical assays for protein characterization

Purified protein concentrations were determined by absorbance at A280 nm. SDS-PAGE was performed under reducing conditions using Novex 12% Tris-Glycine Midi Gels (Thermo Fisher Scientific, Waltham, MA). Separation was performed at 125V for 90 minutes. Gels were stained using Instant Blue Protein Stain (Abcam Inc, United Kingdom), and destained with deionized water for a total of three washes prior to imaging.

### Size exclusion chromatography

Purified RBD protein was separated using a Superose™ 6 Increase 10/300 GL column (Cytiva Life Sciences, catalog no. 29091596) on an ÄKTA Pure 25-L FPLC system (Cytiva Life Sciences, catalog no. 29018224). The column was equilibrated with 3 CVs of a buffer composed of 50 mM sodium phosphate and 150 mM NaCl (pH 7.4) at a rate of 0.25 ml/min. Approximately 1000µg of protein diluted to 500µL with buffer was injected onto the column. One CV of buffer was applied to the column at a rate of 0.25 ml/min for sample elution, and fractions of 0.5mL were collected. All fractions corresponding to each peak were pooled together, concentrated using Pierce™ Protein Concentrator PES, 10K MWCO, 2-6 mL (Thermo Scientific, catalog no. 88516), and the final protein concentration was measured via A280 absorbance.

### Mass spectrometry

Approximately 40-80ug of total protein was digested with PNGaseF, glycerol-free (New England BioLabs, catalog no. P0705L) according to the manufacture’s recommended protocol. Intact mass analysis was performed on a 6530B quadrupole time-of-flight liquid chromatograph mass spectrometer (LC-MS) equipped with a dual ESI source and a 1290 series HPLC (Agilent Technologies, Santa Clara, CA). Mobile phase A consisted of LCMS grade water with 0.1% formic acid, and mobile phase B was LCMS grade acetonitrile with 0.1% formic acid. About 1.0 μg of protein for each sample was injected, bound to a ZORBAX RRHD 300SB-C3 column (2.1 mm × 50 mm, 300 Å, 1.8 μm) (Agilent Technologies, Santa Clara, CA), desalted, and subjected to electrospray ionization. The LC gradient comprised 5% to 95% B over 4 min at a flow rate of 0.8 mL/min. A blank injection using the same LC method between each sample was performed as a wash step. The electrospray ionization parameters were as follows: 350 °C drying gas temperature, 10 L/min drying gas flow, 30 psig nebulizer, 4,000 V capillary voltage, and 250 V fragmentor voltage. Mass spectra were collected in MS mode (0 V collision energy) from 500 to 3,200 m/z at a scan rate of 1 spectra/s. MS spectra were processed using MassHunter BioConfirm software (vB.10.0, Agilent Technologies) using the Maximum Entropy deconvolution algorithm, a mass step of 1 Da, and a mass range of within 50,000 amu appropriate to each protein sample analyzed. For quantitative analysis of the identified P[4]-HMW protein species in the summed mass spectra, area under the curve for each deconvoluted mass peak was used for percent calculations relative to total P[4]-related mass peaks.

## Author’s contributions

N.C.D., C.A.N., S.A.R., and J.C.L. conceived and planned experiments. N.C.D., C.A.N., and R.S.J. generated and characterized yeast strains. S.R.A. performed HPLC assays and protein purifications. C.A.N. performed PNGase treatment and mass spectrometry. N.C.D., C.A.N., S.A.R., and J.C.L. wrote the manuscript.

**Fig. S1.**
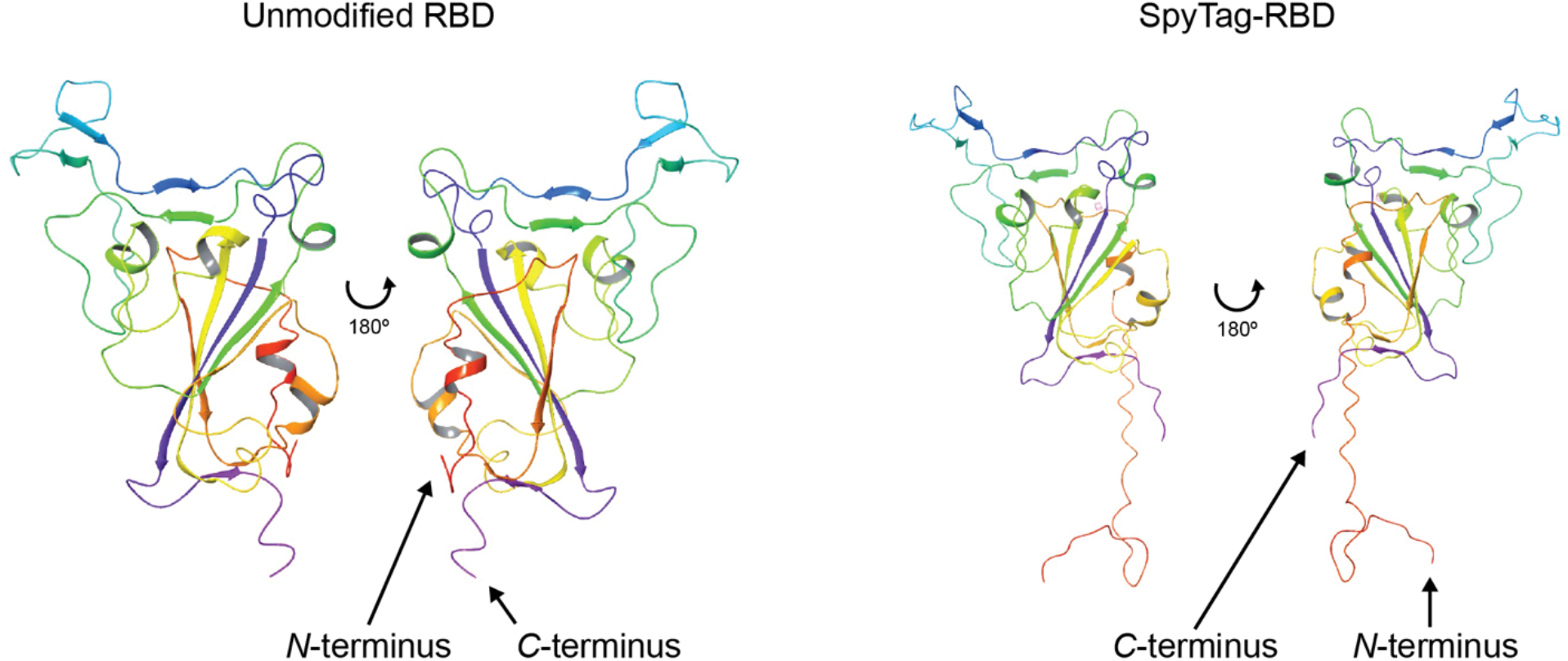
Structural rendering of unmodified RBD and SpyTag-RBD.

